# The Effect of Synthetic Training Data on the Performance of a Deep Learning Based Markerless Biomechanics System

**DOI:** 10.1101/2023.10.19.562758

**Authors:** Ty Templin, Travis Eliason, Omar Medjaouri, David Chambers, Kase Saylor, Daniel P. Nicolella

## Abstract

As markerless motion capture technologies develop and mature, there is an increasing demand from the biomechanics community to provide kinematic data with the same level of accuracy as current gold standard methods. The purpose of this study was to evaluate how adding synthetic data to the training dataset of a deep learning based markerless biomechanics system impacts the accuracy of kinematic measurements during two functional movements. Synchronized video from multiple camera views was captured along with marker-based data from 9 subjects who performed 3 repetitions of countermovement jumps and squats. Including synthetic data to the training reduced lower limb error on average by 65.1% and 70.1% for the countermovement jump and squat movements, respectively. These results demonstrate the promising utility of supplementing the training of a deep learning markerless motion capture system with synthetic data.

## 1 INTRODUCTION

Analysis of human movement has provided critical insight across a broad range of health, disease, and performance related applications. However, the accurate and reliable measurement of movement traditionally requires a dedicated laboratory space with advanced instrumentation operated by highly trained individuals, and is exceedingly time consuming, all of which significantly limit its use in routine clinical and operational assessments. To address the limitations of marker-based motion capture, several markerless motion capture technologies have been developed over the last few decades. Recently, with advances in computer vision and artificial intelligence, deep learning neural network-based markerless systems have begun to demonstrate promising accuracy when compared to marker-based tracking (Kanko et al. 2021; Nakano et al. 2020; Needham et al. 2022).

Neural network based markerless systems require training images and labels, which are typically provided through hand labeled public datasets such as: COCO (Lin et al. 2014), MPII (Andriluka et al. 2014), and Leeds Sports (LSPe, Johnson and Everingham 2010), or proprietary datasets (Kanko et al. 2021). Hand labeling the large number of images required to train a neural network is manually intensive and potentially error prone. In addition, many of these datasets have sparse keypoint labels that limit their utility in 3D kinematic analysis (Needham et al. 2022).

One alternate method that can be used to generate training labels for a markerless system is by projecting 3D labels from optical motion capture to automatically label images from motion capture (mocap) sessions as done in: HumanEva (Sigal, Balan, and Black 2010), Human3.6M (Ionescu et al. 2013), and TotalCapture (Trumble et al. 2017). This technique allows for the automatic labeling of additional points (based on the number of markers in the markerset) and more consistent labeling for occluded points. However, there are some limitations to this type of training data as well. Mocap datasets typically exhibit less background, person, and movement diversity relative to public datasets, which limit the generalizability of systems trained using this data, as well as inherent error due to marker placement inaccuracy, soft tissue artifact, and the inability to directly measure the location of joint centers.

The need for a diverse set of training images that can be automatically labeled in large quantities with high accuracy and fidelity to further improve human pose estimation has led to the creation of synthetic datasets such as AGORA (Patel et al. 2021), SURREAL (Varol et al. 2017), and infiniteform (Weitz et al. 2021). These studies have demonstrated that incorporating these datasets into neural network training results in reduced error in 2D and 3D joint position measurements. However, it remains uncertain how beneficial these datasets are for evaluating more commonly used metrics in biomechanics applications such as joint angles.

Therefore, the purpose of this study is to evaluate the accuracy of the kinematics generated from a markerless biomechanics system (ENABLE) trained with and without synthetic data relative to those of a marker-based system for two functional movements.

## 2. METHODS

### 2.1 Participants and Protocol

8 male subjects aged 20-56 provided informed consent to participate in this study according to an IRB protocol approved by Southwest Research Institute. In this protocol, each subject performed 3 repetitions of countermovement jumps (CMJ) and squats. A video-based markerless system along with a Vicon marker-based motion capture system were used to capture the movement data. We spatially calibrated the global 3D reference frame for the markerless video system such that it coincided with the marker-based global 3D reference frame using a custom procedure and software tool (Eliason et al. 2019). In addition, we determined the camera intrinsic parameters such as the focal length, image center, and distortion coefficients using standard procedures from OpenCV (Bradski 2020). We synchronized the markerless videos in time by using existing synchronization protocols and custom developed signal generation hardware to simultaneously trigger and synchronize data capture and storage from each system.

### 2.2 Markerless Motion Capture

8 FLIR Blackfly cameras recorded video at a frequency of 50 Hz. For the ENABLE markerless analysis, videos were processed through two separate convolutional neural networks (Base and Base + Synthetic) that estimated the location of 49 anatomical points on the human body within each camera view. The core architecture of the two neural networks remained constant (Sun et al., 2019). The only difference between the two networks was the data that was used during training.

For both networks, after the keypoints are detected in each video frame of each of the 8 cameras, a triangulation process is used to estimate the 3D position of each anatomical location. In this process, the calibration information is used to create a set of rays going from the cameras through each of the 2D predictions. A RANSAC (Fischler and Bolles 1981) approach is used to determine a set of inlier and outlier rays and then the 3D point is determined as the point closest to the set of lines using a least squares approach. These 3D locations were then used to scale and drive the kinematics of a musculoskeletal model in OpenSim (version 4.4, Delp et al. 2007). The Rajagopal model (Rajagopal et al. 2016) was modified such that it has 14 degrees of freedom in the lower body: 3 hip, 3 knee, 2 ankles. Ankle subtalar angle includes both transverse and frontal plane motion but for reporting purposes will be included in the ab/adduction columns.

This model was scaled to match each subject’s anthropometry by using median segment lengths of eachsegment during each trial. After the model was scaled, an inverse kinematic fit globally optimized the pose of the model in each frame using the python API interface with the OpenSim Inverse Kinematics (IK) tool.

### 2.3 Base Condition

The Base neural network was trained on approximately 100,000 images from two broad source classes: public datasets (∼50,000 images) and auto labeled images from a marker-based system (∼50,000 images). The public images consisted of select images from the COCO foot, MPII, and LSPe datasets. The MPII and LSPe training sets are labeled with 15 points including joints centers at the hip, knee, ankle, shoulder, elbow, wrist, and head keypoints. The COCO foot dataset includes the same labels as well as additional labels of the foot keypoints (Cao et al. 2018). The optical motion capture dataset consisted of images that were collected in a motion capture lab (separate from where the current study was conducted) of individuals performing a variety of functional and sports related movements. These images were labeled using a patented approach where 49 3D anatomical locations were determine based on the model pose from a marker-based approach that were reprojected into the 2D image space (Saylor et al. 2019). The 49 keypoints consisted of the same 25 keypoints utilized in the COCO dataset plus additional keypoints throughout the body to ensure that each of the following body segments had a minimum of three keypoints attached to them: head, torso, upper arms, forearms, hands, thighs, shanks, and feet (Figure 1). This allowed for full 6 degree of freedom (DOF) characterization of each segment. The keypoints not included in the public images were ignored during the training.

**Figure 1.**
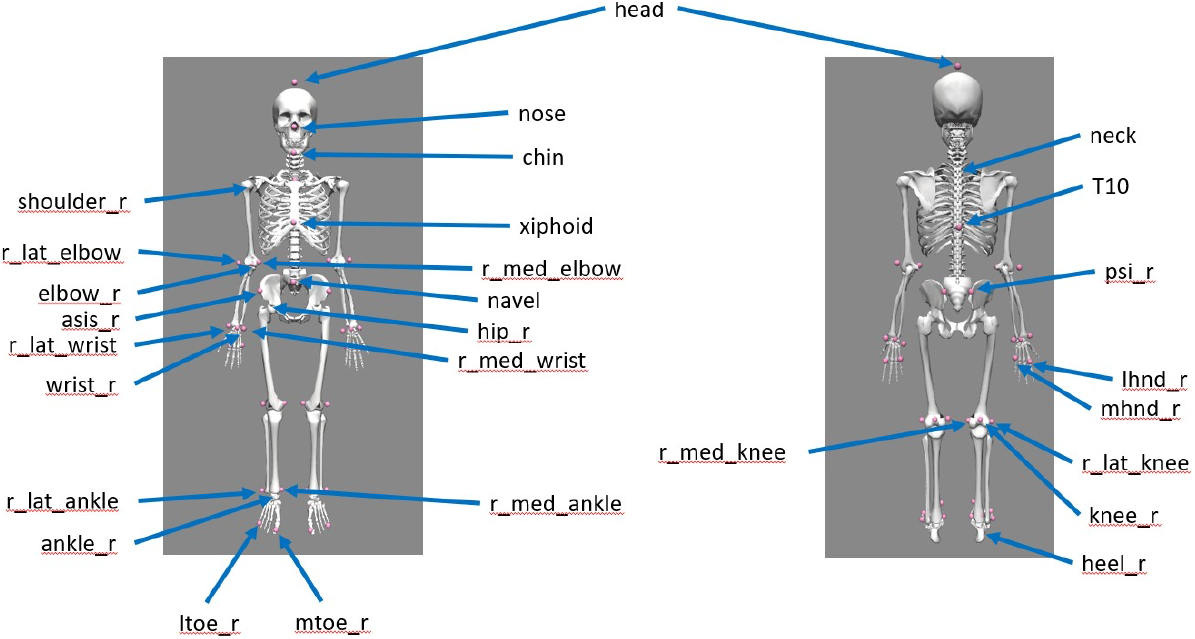
Keypoint annotations on OpenSim model. Markers that are designated as being on the right side of the body also have a corresponding point on the left side of the body.

### 2.4 Base + Synthetic Condition

The Base + Synthetic neural network was trained on the same datasets described above plus a synthetic dataset generated using a custom application of the Infinity API (Infinity). This dataset (∼50,000 images) consisted of synthetic people performing squats and countermovement jumps with varying body size, shape, camera position, and appearance in backgrounds that mimicked the present data collection background and was annotated using the same 49 keypoints as the optical motion capture dataset.

### 2.5 Marker-based Motion Capture

The optical marker-based motion capture was collected using a 16 camera Vicon system at 100Hz. A full body markerset was used with 84 markers. 36 of these markers were placed on bony landmarks. The remaining 48 markers were attached as 4 marker clusters to the following segments bilaterally: thigh, shank, foot, upper arm forearm, hand. Before performing any of the movements, a static trial was captured with the subject in a T-pose. This capture was used to scale the same OpenSim model for each individual as described for the markerless captures. After the static trial was collected, the medial and lateral markers of the elbow, wrist, knee, ankle, and toe markers were removed. After collecting the movement trials, the marker data was filtered with a 4^th^ order bi-directional butterworth filter with a 6Hz low pass cutoff frequency. This filtered data was then used as input to drive inverse kinematics (IK) for the scaled model.

### 2.6 Data Analysis

To evaluate the performance of the ENABLE markerless system trained with and without synthetically produced training data, the IK results for the hip, knee, and ankle were compared to the IK results determined from the marker-based system. The comparison was made using root mean squared error (RMSE) and Pearson Correlations.

## 3 RESULTS

The Base condition (Figure 2) produced results that were generally noisier with greater deviation from Vicon than the Base + Synthetic condition (Figure 3). The largest difference between the base and base + synthetic condition occurred at ankle flexion. The base condition had poor tracking of the ankle with >44 degrees RMSE for both the squat and CMJ with Pearson correlation of <0.34, while the base + synthetic condition produced ankle flexion results much more like the Vicon results with RMSE <8 degrees and correlations >0.96 (Table 1). Each of the other DOFs also showed reduced RMSE and greater Pearson correlation relative to the Vicon data. In the Base + Synthetic condition, the largest error was observed at the subtalar ankle angle with RMSE >12 degrees and correlations <0.3. All other joint angles showed RMSE <9 degrees. Hip internal/external rotation and ad/abduction had moderate correlations ranging from 0.48 - 0.75 and were higher for the squat than for the CMJ. Knee internal/external rotation and ad/abduction had lower correlation ranging from 0.11 - 0.58 with higher correlations observed during the CMJ compared to the squat.

**Figure 2.**
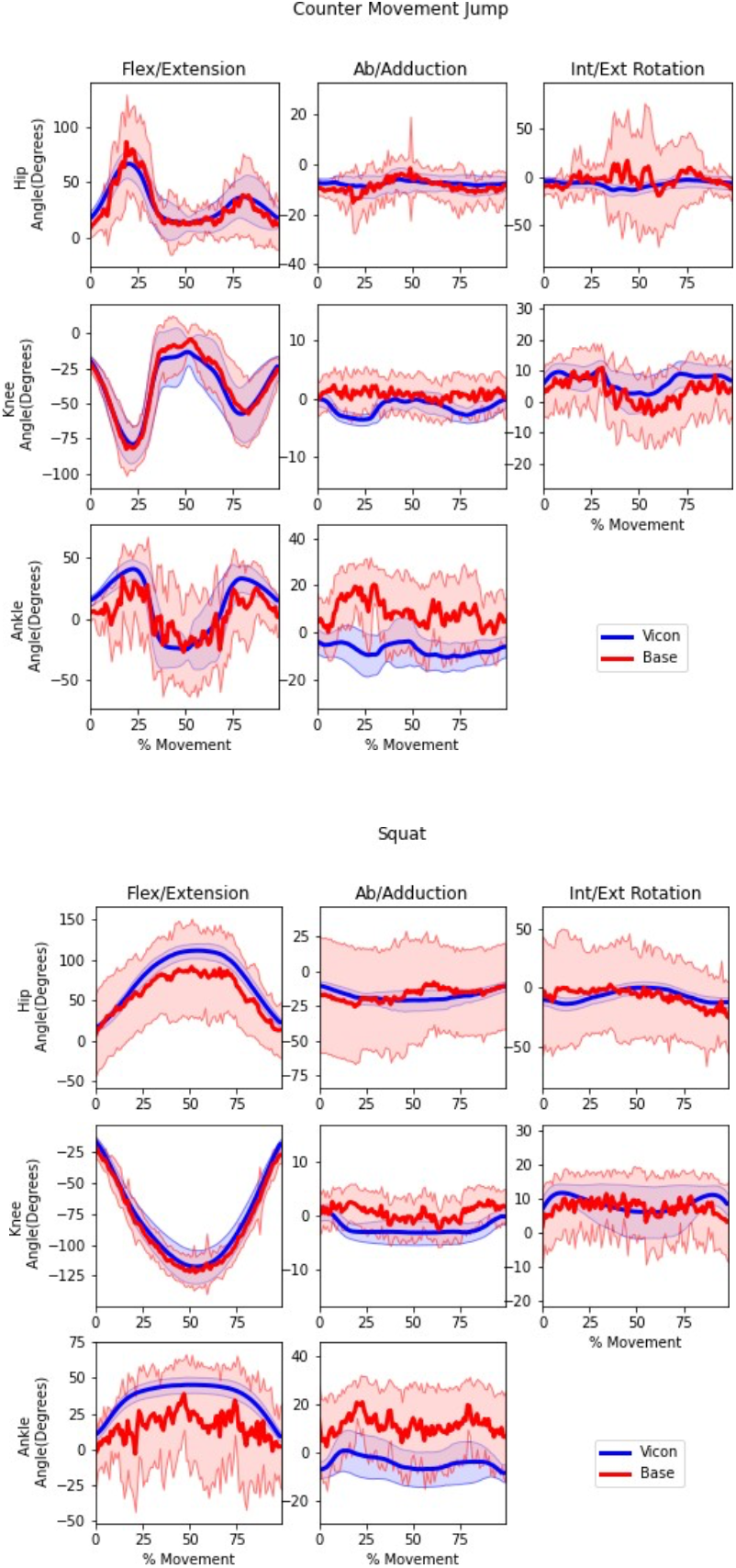
Mean +/-1 standard deviation of the time history trajectories of lower limb kinematics measured by Vicon (blue) and the Base markerless system (red) during countermovement jumps and squats.

**Figure 3.**
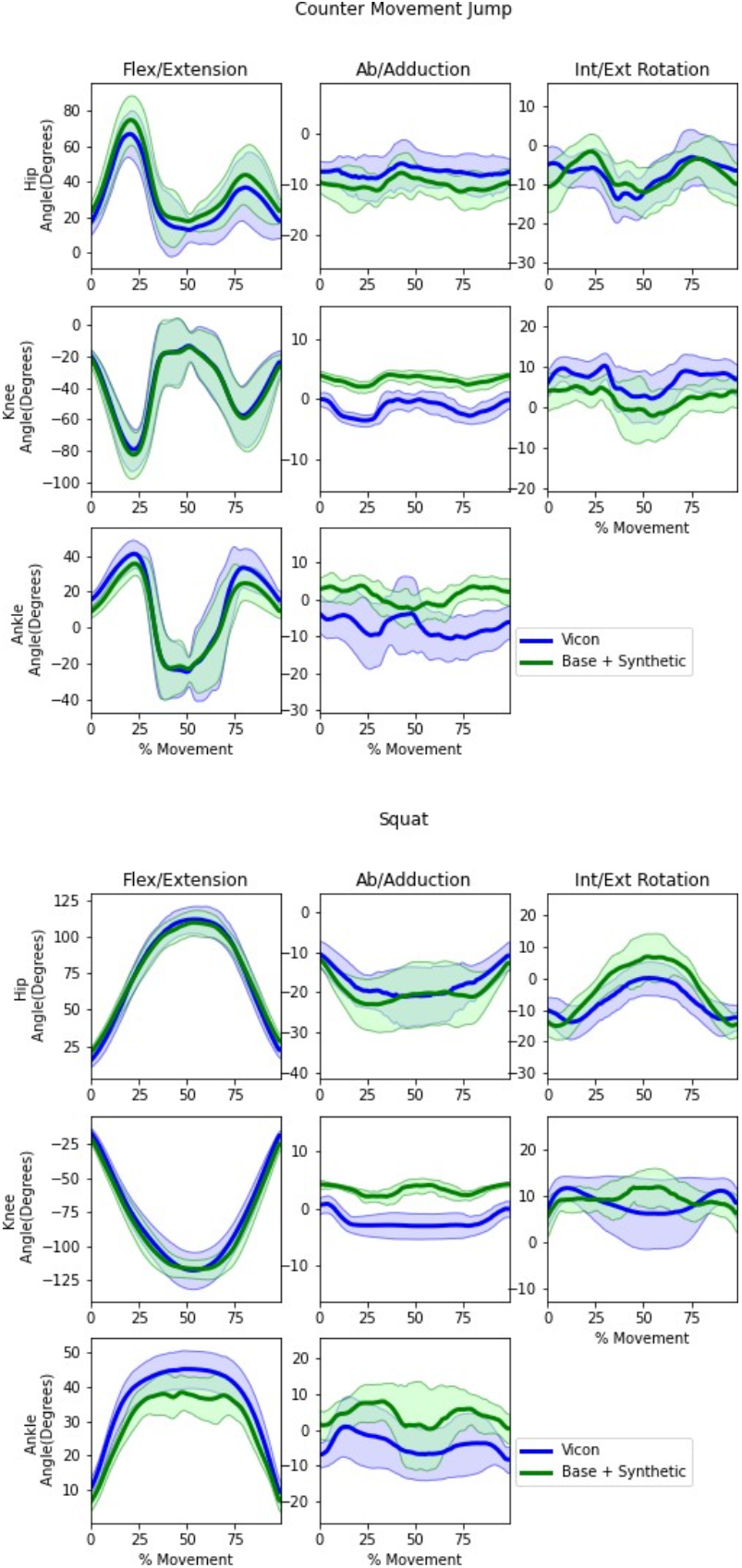
Mean +/-1 standard deviation of the time history trajectories of lower limb kinematics measured by Vicon (blue) and the Base + Synthetic markerless system (green) during countermovement jumps and squats.

**Table 1.**
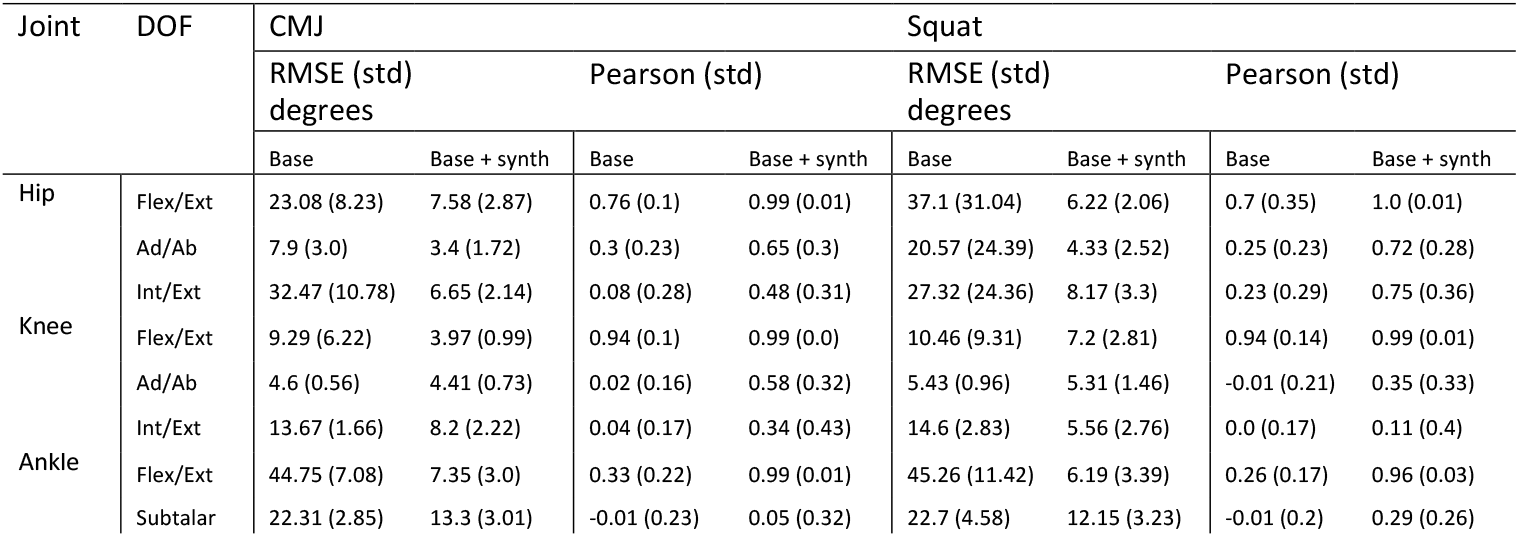
Comparison of RMSE +/-1 standard deviation and Pearson correlation +/-1 standard deviation between both markerless systems relative to Vicon for lower limb kinematics of countermovement jumps and squats.

## 4 DISCUSSION

This study aimed to evaluate the accuracy of the ENABLE markerless motion capture system trained with and without synthetic training images. The average lower limb error was 65.1% and 70.1% lower in the Base + Synthetic condition than the Base condition relative to the Vicon measurements for the CMJ and Squat movements respectively (Figure 4).

**Figure 4.**
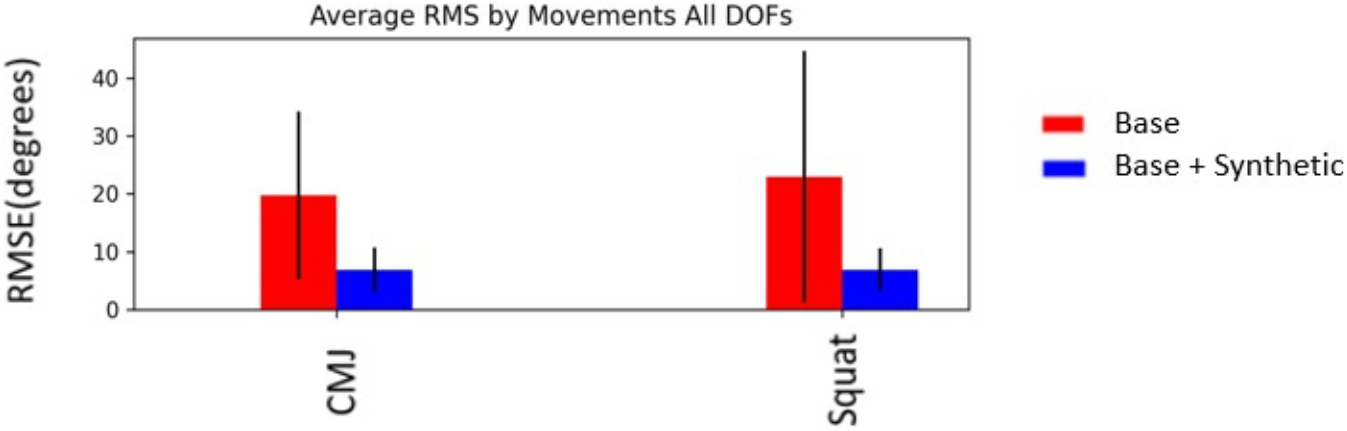
Comparison of Average RMSE for Lower Limb Kinematics between Base (red) and Base+Syntethic Markerless Conditions (blue) for CMJ and Squat movements.

Knee flexion angles were not substantially different between the two conditions compared to the Vicon results. The hip flexion results shown in the present study are more accurate than results from a previous study which report errors of >22 degrees for squats (Ito et al., 2022) and >16 degrees for countermovement jumps (Strutzenberger et al. 2021). One possible explanation for this finding is that the markerless system used in this study identifies points on the pelvis, which many markerless systems are unable to track, due to the sparse labels of many commonly used datasets. The Base + Synthetic trained condition showed largest improvements relative to the base condition in the ankle non-sagittal plane DOFs. These DOFs in the Base condition exhibited greater error than what has previously been reported in the literature for other markerless systems as these DOFs were often noisy and differed from trial to trial. This finding in the Base condition is likely a result of the lack of adequate diversity in the training data that contained all 49 points. In this condition, the only dataset that contained labels for each point was the data collected in the motion capture lab. The variability in this dataset was limited because it included one background and 15 subjects. Therefore, the base network had difficulty generalizing the prediction of the additional points for different subjects and backgrounds. However, the Base + Synthetic condition produced much more consistent and reliable predictions for the ankle flexion and non-sagittal DOFs. The additional variability inherent in the synthetic data in categories such as clothing type, body shape, background, movement as well as the consistently pixel perfect labels improved the predictions of those points.

Another potential application for synthetic data usage is for specific use cases in environments, camera orientations, movement patterns that are underrepresented in public datasets. In these applications, synthetic images could be generated to mimic the expected environment to help improve robustness and accuracy. The ability to generate numerous images with accurate labels makes synthetic data a promising solution over the time consuming and potentially error prone and inconsistent alternative of hand labeling. In addition, the flexibility in the location and number of points afforded by synthetic data provides the potential to generate more detailed representations of specific degrees of freedom that are difficult to resolve using the sparse keypoints associated with most training datasets.

While synthetic data improved the results in the frontal and transverse plane at the knee and ankle, these DOFs still exhibited the greatest discrepancy in correlation and RMS to the marker data. One potential cause of the persisting error in the transverse and frontal planes is that the off-axis points used to define the femur and tibia segments were directly medial and lateral of the knee and ankle joint centers. The relative proximity of these points to each other may cause any error in the predicted 3D position of the point to magnify the error during IK. These off-axis points were chosen as labels in the synthetic data to reflect traditional marker-based marker sets typically located on a subject using anatomical landmarks. However, synthetic data does not require keypoints to be located on easily accessible anatomical locations (e.g., bony landmarks). Therefore, work should be done to investigate the benefit of placing non-sagittal keypoints in a more optimal locations on each segment to reduce the impact of error on the IK results.

Another potential cause of the decreased accuracy in non-sagittal planes of motion at the knee and ankle is error propagation to distal segments of the kinematics tree. Since the IK solver used in OpenSim is a global optimization procedure, the error observed in the joint may be compounded by error in joints that are more proximal to the root joint (pelvis). Finally, the kinematic structure of the synthetic models only includes flexion and extension of the knee and ankle and does not include ab/adduction or internal/external rotation. In the future, these DOFs should be added to the synthetic body model to replicate the desired movements more accurately. However, since the neural network is trained to identify features in the images and does not directly predict joint angles, it is likely that this limitation may be less important to the overall performance of the system.

While this study used a marker-based system as the ground truth, there are well documented limitations of marker-based systems that can result in erroneous measurements. These include skin artifacts and lack of consistent marker placement (Benoit, Damsgaard, and Andersen 2015; Benoit et al., 2006; Cappozzo et al., 1996; Fuller et al., 1997; Gorton et al., 2009; Kessler et al. 2019; Leardini et al., 2005; Lucchetti et al., 1998; Miranda et al. 2013; Peters et al., 2010; Reinschmidt et al., 1997). In addition, the marker-based systems also require substantial manual effort to post process and gap fill missing marker trajectories. Analogous manual editing of the markerless points was not included as part of this research but could be an added feature for users who are experienced with this data post processing method. Despite these limitations of marker-based tracking, it is considered the standard method of acquiring accurate kinematic information in the biomechanics community due to the difficultly of collecting data using the gold standard biplane fluoroscopy equipment.

The results of this study highlight the promising potential of using synthetic data to augment training of a deep learning based markerless motion capture system. In this study, we found that including synthetic data into the training of the ENABLE markerless system substantially improved of the accuracy and reliability for two functional movements. Further investigation into the benefit of using synthetic training data in other environments to fine-tune markerless motion capture system performance is warranted.

